# Mosquito collection method assessments for xenomonitoring in two cross-bordering villages with different ecosystems, Yanfolila, Mali, 2022-2023

**DOI:** 10.64898/2026.06.22.733916

**Authors:** Lamine Soumaoro, Michel Emmanuel Coulibaly, Abdallah Amadou Diallo, Moussa Sangaré, Abdoul Fatah Diabaté, Siaka Yamoussa Coulibaly, Salif Seriba Doumbia, Mahamoud Mahamadou Koureichi, Abdoul Karim Koné, Karim Dembélé, Housseini Dolo, Alpha Seydou Yaro, Yaya Ibrahim Coulibaly

**Affiliations:** Neglected Tropical Diseases Research and Training Unit, Infectious Diseases and Medical Entomology Research and Training Center (IDMERTC), University of Sciences, Techniques and Technologies of Bamako (USTTB), Bamako, Mali; Faculty of Sciences and Techniques (FST), University of Sciences, Techniques and Technologies of Bamako (USTTB), Bamako, Mali

**Keywords:** Xenomonitoring, lymphatic filariasis, collection methods, *Anopheles gambiae* complex, *Culex* spp., Konfra and Siradjouba

## Abstract

Entomological surveillance is essential for the control of lymphatic filariasis (LF). This xenomonitoring study, conducted in 2022–2023, evaluated the effectiveness of three mosquito trapping methods to identify the most suitable tool for surveillance following the discontinuation of mass drug administration. The surveys took place in two Malian villages: Konfra (seasonal watercourse) and Siradjouba (permanent watercourse). Three techniques were compared: Pyrethrum spray catch (PSC) conducted at the end of each visit inside 30 randomly selected houses, and different collection points were chosen based on the characteristics of the study area to place the Ifakara Type C tent (Ifakara) and the gravid trap (GT). Collections were conducted simultaneously at both sites. All *Anopheles gambiae* and *Culex* spp. collected in 2022–2023 were morphologically identified and then analyzed in the laboratory for their infectious status. Data was processed using SPSS v25; Fisher’s exact test was used to compare proportions, with a significance threshold of p < 0.05. A total of 4,732 mosquitoes were captured (95.05% in 2022). In 2022, the *Anopheles* were predominant in Konfra (91.38%), while *Culex* predominated in Siradjouba (63.41%). In 2023, *Anopheles* was found only in Konfra (3 specimens), while *Culex* was predominant in Siradjouba (64.07%). In 2022, PSC collected 90.57% of *Anopheles*, Ifakara 9.43%, and GT 0%. For *Culex*, GT collected 53.67%, PSC 36.62%, and Ifakara 9.71%. In 2023, the few *Anopheles* (3) came from PSC; *Culex* were mainly collected by the Ifakara tent (43.29%). For FL xenomonitoring, the combination of PSC and the GT appear to be effective collection methods.

## Introduction

Lymphatic filariasis (LF), a debilitating parasitic disease commonly known as elephantiasis, is caused by nematodes such as *Wuchereria bancrofti* (the main species in West Africa), *Brugia malayi*, and *Brugia timori* [1–3]. Primarily transmitted by the *Anopheles gambiae* complex in West Africa, although *Mansonia* is implicated in Guinea and Ghana, it remains a major challenge despite efforts by the Global Programme to Eliminate Lymphatic Filariasis (GPELF) through mass drug administration (MDA) campaigns [1,2]. Post-MDA entomological surveillance is essential for detecting residual transmission, particularly in areas with low vector densities, where collection tools must target mosquitoes with varied behaviors (endophilic, exophilic, anthropophilic, zoophilic) according to ecological contexts and objectives [4,5].

Xenomonitoring, recommended by the GPELF, assesses the presence of filarial larvae in mosquitoes via dissection or, preferably, qPCR for sensitive and non-invasive detection of pathogen DNA/RNA [2,6,7] This approach complements human blood analysis and also evaluates the effectiveness of programs against other vector-borne diseases [8]. However, the efficacy of collection methods (e.g., pyrethrum spray catch, tent traps, gravid traps) varies by target species and ecosystems, necessitating tailored tools for optimal post-MDA surveillance [9,10].

In Mali and Guinea, neighboring countries where LF is endemic, Mali conducted its third transmission assessment survey (TAS3) in 2019 in the Bougouni-Yanfolila evaluation unit, while no such assessment had been performed in Guinea at that time. Border areas, characterized by intense human exchanges, pose a high risk of LF reemergence in Yanfolila. As Mali progresses toward LF elimination, validated tools for post-elimination surveillance are crucial, especially in villages with contrasting ecological profiles like Siradjouba (permanent watercourse) and Konfra (temporary watercourse) in southern Mali [11].

This study compares the effectiveness of three mosquito capture methods, namely pyrethrum spray catch (PSC), the Ifakara Type C tent (Ifakara), and the gravid female trap (GT), during a xenomonitoring study conducted in 2022-2023 in these two border villages [11]. The main objective is to identify the most performant method in terms of mosquitoes captured, to select the optimal tool for LF xenomonitoring in Mali and optimize its recent integration into post-MDA activities.

## Methodology

### Study sites

The study was carried out in two villages (Siradjouba and Konfra) bordering the Yanfolila health district, located in the Sikasso region, southern Mali (figure 1). This area is characterized by an intense cross-border mobility, making it a priority site for LF surveillance after MDA [12]. Siradjouba and Konfra present contrasting hydrological ecologies, enabling a comparative assessment of vector densities and the performance of mosquito collection methods in different settings. Siradjouba in the Kabaya health sub-district is crossed by a permanent watercourse, while Konfra in the Nièssoumala health sub-district has a temporary watercourse.

**Figure 1:**
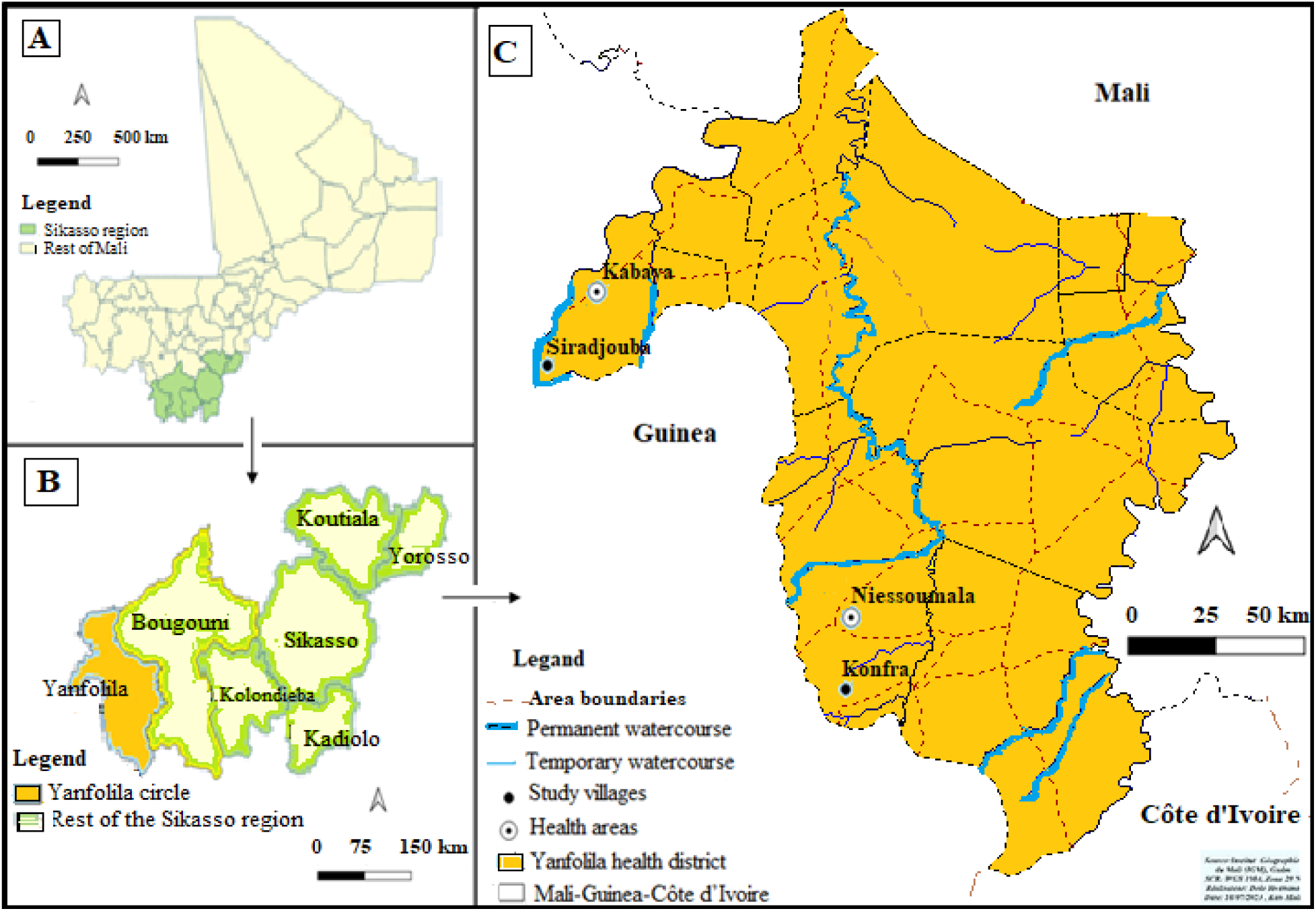
Map showing the two study villages in the Yanfolila health district. A: Map of Mali showing the targeted border area. B: Map showing the Sikasso region with the boundaries of Yanfolila health district. C: Map of the Yanfolila health district showing the sampling sites.

### Types and periods of study

This is a cross-sectional study, based on quantitative entomological data collection. Three mosquitoes collection sessions were organized, in June 2022, August 2022, and January 2023. Each session consisted in simultaneous implementation of three collection methods in the two study villages for 10 days.

### Collection methods

In both villages, three techniques for collecting adult mosquitoes were implemented simultaneously in 2022 and 2023: the Ifakara Type C tent (Ifakara), the gravid trap (GT), and the Pyrethrum Spray Catch (PSC) [13–15]. The locations for the Ifakara tent and GT were selected based on the characteristics of the study area. The PSC was deployed at the end of each collection session in 30 randomly selected inhabited huts, with the support of village representatives. The specimens collected from the Anopheles gambiae complex and the *Culex spp*. complex were identified morphologically on site.

### Data collection

Each collection session consisted of ten consecutive days of operations. In each village, two GTs and two Ifakara tents were deployed simultaneously, operating from 6 p.m. to 6 a.m. the following day in order to target the mosquitoes’ nocturnal activity. These traps were set up in separate ecological zones within each of the two villages for nine consecutive days. The 10th day was dedicated to a single PSC session in each village studied. Community representatives selected 30 randomly, and collections were carried out from 8 am to 6 pm.

After each collection session, the insects were sorted by trap, village, genus (*Anopheles*, *Culex*), and others. Only *Anopheles* and *Culex* complexes members were considered in this study. The collected mosquitoes were stored by collection method, room, genus, and pool. They were kept in 1.5 ml Eppendorf tubes containing 70% ethanol. The tubes were labeled with an identification number made with numbers related to the health district, village, collection method, and collection. The geographic coordinates of the trap installation point, and mosquito collection rooms were systematically recorded. Entomological data were recorded on printed sheets on fields before being entered into a Microsoft Excel file. The collected mosquitoes were preserved in 70% ethanol in the field at room temperature. They were grouped by trap, hut, and village on site. They were then sent to the laboratory, stored at −20°C, and divided into batches of 20 individuals.

### Sample processing in the laboratory

Collected mosquitoes were processed in the laboratory to detect *Wuchereria bancrofti* DNA by real-time qPCR protocol, using the StepOne Plus™ machine from Applied Biosystems [11].

### Data analysis and statistical considerations

Fisher’s exact test was used to compare mosquito proportions with 95% confidence intervals. The following entomological parameters were determined: vector fauna composition, vector densities.

### Ethical considerations

The Ethics Committee of the University of Sciences, Techniques and Technologies of Bamako approved the protocol under number 2022/294/CE/USTTB.

## Results

This study evaluates mosquito collection methods (PSC: indoor insecticide spraying; Ifakara: human bait trap; and GT: gravid female trap) for xenomonitoring in two villages in Yanfolila (Mali): Konfra (temporary watercourses) and Siradjouba (permanent watercourses). It focuses on the *Anopheles gambiae* complex (the main vector for malaria and lymphatic filariasis) and *Culex spp*. (vectors for arboviruses). The results highlight marked differences by village, gender, year, and method.

### 1. Specific composition and geographical distribution of mosquitoes collected in the two villages of the Yanfolila health district in 2022 and 2023

A total of 4,732 specimens of both mosquito genera were collected in the villages of Konfra and Siradjouba between 2022 and 2023, of which 95.05% (4,498/4,732) were captured in 2022 and 4.95% (234/4,732) in 2023.

Of the 4,498 mosquitoes collected in 2022, the *Anopheles* complex accounted for 21.92% (986/4,498) and the *Culex spp*. genus accounted for 78.08% (3,512/4,498). Of the 986 *Anopheles*, the majority were captured in Konfra (91.38%; 901/986; 95% CI: [89.45-93.06]), while 8.62% were collected in Siradjouba. For the 3,512 *Culex spp*., the situation was reversed: this genus was most abundant in Siradjouba (63.41%; 2,227/3,512; 95% CI: [61.79-65.01]) and was least represented in Konfra (36.59%; 1,285/3,512; 95% CI: [34.99-38.21]) (Table 1).

**Table 1:** Species composition and geographic distribution of mosquitoes collected in the two study villages in 2022-2023.

| Year | Specimens | Konfra (n; %) | [95% CI] | Siradjouba (n; %) | [95% CI] | Total (%) | [95% CI] |
| --- | --- | --- | --- | --- | --- | --- | --- |
| 2022 | *Anopheles gambiae* complex | 901; 91.38% | [89.45-93.06] | 85; 8.62% | [6.94-10.55] | 986 (21.92) | [20.72-23.16] |
|  | *Culex spp.* | 1 285; 36.59% | [34.99-38.21] | 2 227; 63.41% | [61.79-65.01] | 3 512 (78.08) | [76.84-79.28] |
|  | Sous-total 2022 |  |  |  |  | 4 498 (100) | [99.92-100] |
| 2023 | *Anopheles gambiae* complex | 3; 100% | [29.24-100] | 0; 0% | [0.0-70.76] | 3 (1.28) | [0.27-3.70] |
|  | *Culex spp.* | 83; 35.93% | [29.74-42.48] | 148; 64.07% | [57.52-70.26] | 231 (98.72) | [96.3-99.73] |
|  | Sous-total 2023 |  |  |  |  | 234 (100) | [98.44-100] |
| Total |  |  |  |  |  | 4 732 |  |

Of the 234 mosquitoes collected in 2023, only 3 specimens of *Anopheles gambiae* complex were represented, i.e., 1.28% (3/234), and the genus *Culex spp*. 98.72% (231/234). The three *Anopheles* were captured in Konfra (100%; 3/3; 95% CI: [29.24-100]), while no *Anopheles* were captured in Siradjouba. Conversely, (64.07%; 148/234; 95% CI: [57.52-70.26]) of *Culex spp*. were captured in Siradjouba and the lowest number in Konfra (35.93%; 83/234; 95% CI: [29.74-42.48]) (Table 1).

### 2. Effectiveness of *Anopheles gambiae* complex capture methods by location and year

A total of 989 specimens of the *Anopheles gambiae* complex were collected using the three capture methods in the two villages, including 986 in 2022 and only 3 in 2023.

Of the 986 individuals collected in 2022, the majority (91.38%; 901/986) came from Konfra, while 8.62% (85/986) were captured in Siradjouba. The PSC method yielded the highest proportion, 90.57% (893/986; 95% CI: [88.57-92.32]), followed by Ifakara with 9.43% (93/986; 95% CI: [7.68-11.43]). No captures were made with the GT in either village in 2022. Of the 893 individuals captured by PSC, 91.04% (813/893; 95% CI: [88.97-92.83]) came from Konfra and 8.96% (80/893; 95% CI: [7.16-11.03]) from Siradjouba. Of the 93 individuals collected with the Ifakara, 94.63% (88/93; 95% CI: [87.90-98.23]) were captured in Konfra and 5.37% (5/93; 95% CI: [1.76-12.10]) in Siradjouba (Table 2).

**Table 2:** Comparison of the number of *Anopheles gambiae* complex captured by locality and collection method.

| Specimens | Villages | Total collected (%) | PSC | [95% CI] | IFA | [95% CI] | GT | [95% CI] |
| --- | --- | --- | --- | --- | --- | --- | --- | --- |
| *Anopheles gambiae* complex 2022 | Konfra | 901 (91.38) | 813 (91.04) | [88.97-92.83] | 88 (94.63) | [87.90-98.23] | 0 (0) | [0-100] |
|  | Siradjouba | 85 (8.62) | 80 (8.96) | [7.16-11.03] | 5 (5.37) | [1.76-12.10] | 0 (0) | [0-100] |
|  | **Total 1** | **986 (100)** | **893 (90.57)** | **[88.57-92.32]** | **93 (9.43)** | **[7.68-11.43]** | **0 (0)** | **[0-0.37]** |
| *Anopheles gambiae* complex 2023 | Konfra | 3 (100) | 3 (100) | [29.24-100] | 0 (0) | [0-100] | 0 (0) | [0-100] |
|  | Siradjouba | 0 (0) | 0 (0) | [0.0-70.76] | 0 (0) | [0-100] | 0 (0) | [0-100] |
|  | **Total 2** | **3 (100)** | **3 (100)** | **[29.24-100]** | **0 (0)** | **[0.0-70.76]** | **0 (0)** | **[0.0-70.76]** |
| **Total (1+2)** |  | **989 (100)** | **896 (90.60)** | **[88.60-92.34]** | **93 (9.40)** | **[7.66-11.40]** | **0 (0)** | **[0.0-70.76]** |

PSC=Pyrethrum spray catch, IFA=Ifakara tent trap, GT=Gravid trap

In 2023, only three specimens of the *Anopheles gambiae* complex were captured, exclusively in Konfra using the PSC method (100%; 3/3; 95% CI: [29.24-100]) (Table 2).

### 3. Effectiveness of *Culex spp*. capture methods by location and year

The three methods collected 3,743 *Culex spp*. specimens in the two villages, including 3,512 in 2022 and 231 in 2023.

For the 3,512 *Culex spp*. individuals collected in 2022, the situation was reversed: this genus was most abundant in Siradjouba (63.41%; 2,227/3,512; 95% CI: [61.79-65.01]) and was the least represented in Konfra (36.59%; 1,285/3,512; 95% CI: [34.99-38.21]). Among the 3,512 *Culex spp*. specimens captured, GT was the most productive with 53.67% (1,885/3,512) of captures (95% CI [52.01-55.33]), and a significantly higher yield in Siradjouba (79.42% (1,497/1,885) of captures, 95% CI [77.52-81.22]) compared to only 20.58% (388/1,885) (95% CI [18.78-22.48]) in Konfra. Among the 3,512 *Culex spp.* specimens captured, PSC collected 1,286 individuals, or 36.62% (1,286/3,512) (95% CI [35.02-38.24]), and it was more effective in Konfra (59.48% (765/1,286); 95% CI [56.75-62.18]) than in Siradjouba (40.52% (521/1,286); 95% CI [37.82-43.25]). Of the 3,512 *Culex spp*. specimens collected, Ifakara captured 9.71% (341/3,512); 95% CI [8.75-10.74]. It performed better in Siradjouba (61.29% (209/341); 95% CI [55.89-66.49]) than in Konfra (38.71% (132/341); 95% CI [33.51-44.11]) (Table 3).

**Table 3:**
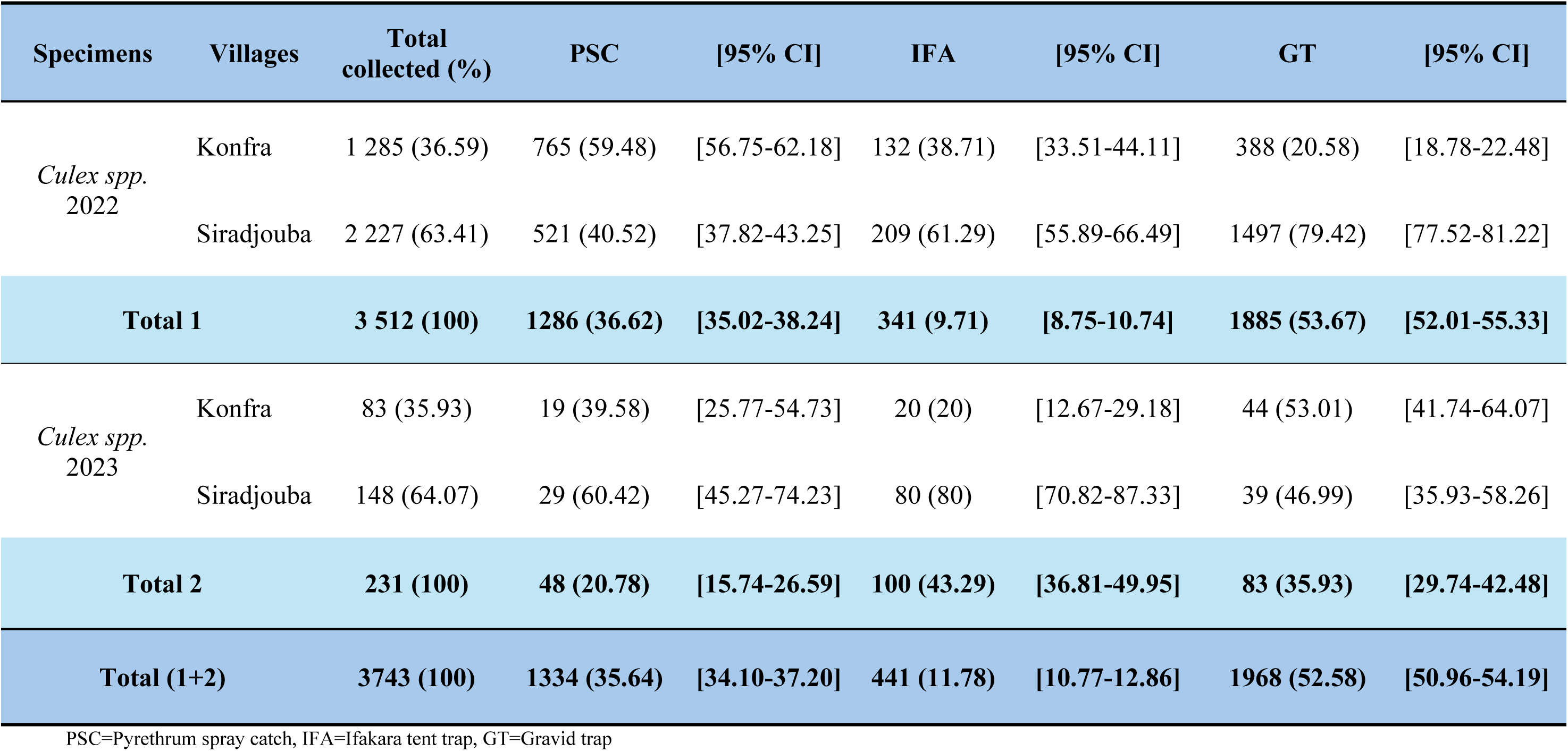
Comparison of the number of *Culex spp.* caught by locality and by collection method.

In 2023, a total of 231 specimens of *Culex spp*. were collected. The largest number was captured with the Ifakara tent, accounting for 43.29% (100/231) with a 95% CI [36.81-49.95], followed by the GT and then the PSC with respectively (35.93% (83/231); 95% CI [29.74-42.48]) and (20.78% (48/231); 95% CI [15.74-26.59]). Among the 100 *Culex spp*. specimens captured with the Ifakara tent, the trap became dominant in Siradjouba 80% (80/100); 95% CI [70.82-87.33]), compared to only 20% (20/100) in Konfra (95% CI [12.67-29.18]). Of the 83 *Culex spp*. specimens captured with the GT, performance was relatively balanced between the two locations, with a slight predominance in Konfra (53.01% (44/83); 95% CI [41.74-64.07]) compared to Siradjouba (46.99% (39/83); 95% CI [35.93-58.26]). Of the 48 *Culex spp*. specimens captured, the PSC performed better in Siradjouba (60.42% (29/48); 95% CI [45.27-74.23]) than in Konfra (39.58% (19/48); 95% CI [25.77-54.73]) (Table 3).

### 4. Change in the total number of mosquitoes captured in the two villages in 2022 and 2023

A clear difference in the species composition of mosquitoes was observed between the two villages. The Konfra site showed a much higher abundance of the *Anopheles gambiae* complex, with 904 specimens collected, compared to only 85 in Siradjouba in 2022 and 2023. In contrast, for *Culex spp*., the trend was reversed, with Siradjouba showing a significantly higher density, with 2,375 individuals collected, compared to 1,368 in Konfra in 2022 and 2023 (Figure 2)

**Figure 2:**
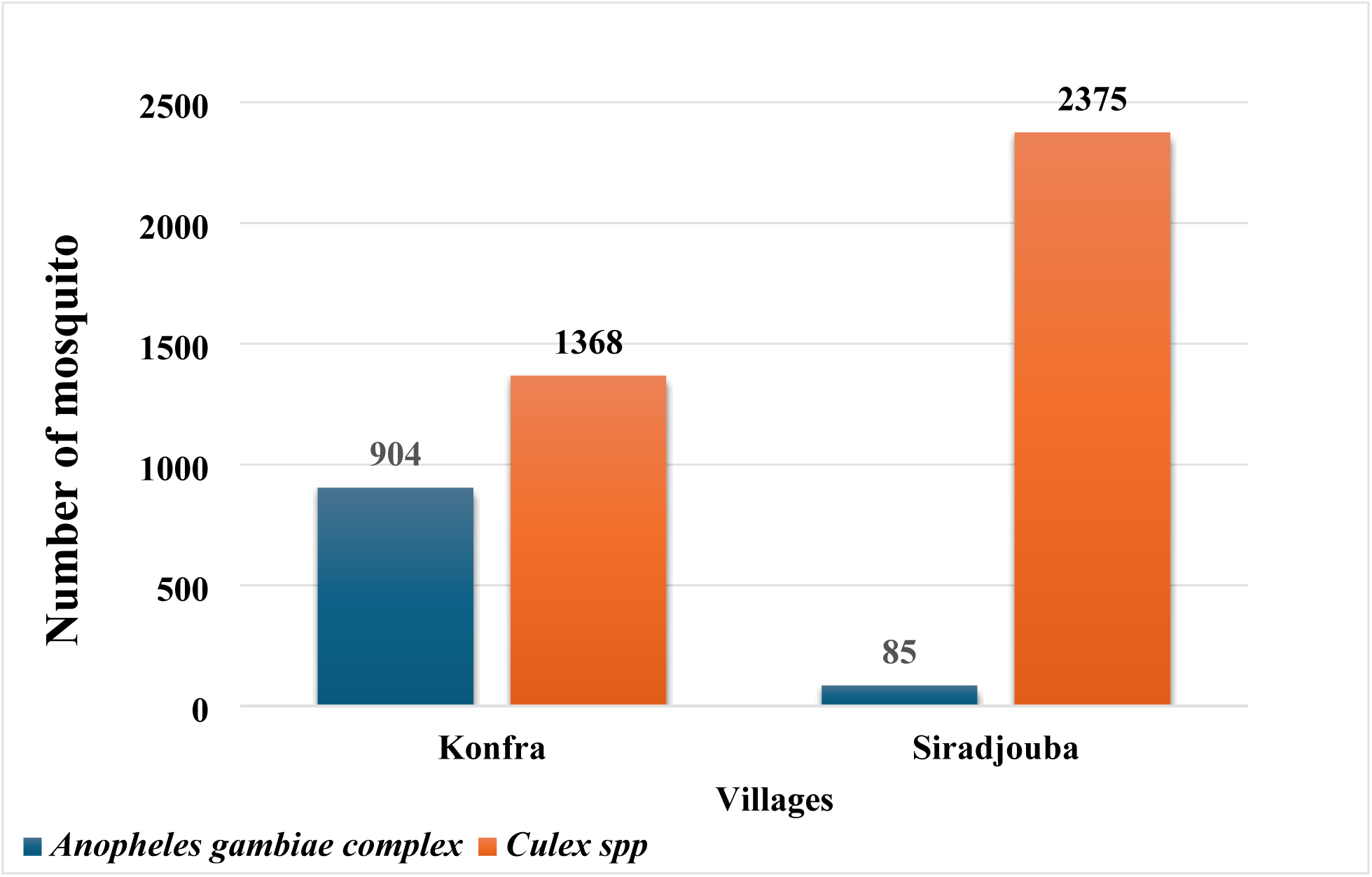
Changes in the total number of mosquitoes caught in the two villages in 2022 and 2023.

## Discussion

Between 2022 and 2023, a total of 4,732 mosquitoes were collected in the villages of Konfra and Siradjouba. In 2022, the *Anopheles gambiae* complex accounted for 20.84% of captures (986 individuals), mainly in Konfra (91.38%), while *Culex spp*. (3,512 individuals) were mainly present in Siradjouba (63.41%). In 2023, the density of *Anopheles gambiae* complex decreased significantly, with only 3 specimens captured, all in Konfra, while *Culex spp*. remained dominant, with a majority in Siradjouba (64.07%). These results highlight the spatial and temporal dynamics of *Anopheles gambiae* complex and *Culex spp*. populations, as well as the comparative effectiveness of the capture methods used in Konfra and Siradjouba in 2022-2023.

### Specific composition and geographical distribution of mosquitoes collected in the two villages of the Yanfolila health district in 2022 and 2023

The results reveal an overall dominance of *Culex spp*. (78% in 2022; 99% in 2023) over the *Anopheles gambiae* complex, with a marked geographical distribution. Thus, the village of Konfra favored *Anopheles* (91% of captures in 2022), while Siradjouba favored *Culex spp*. (63% in 2022). These trends can be explained by local ecological factors: in Konfra, the proximity of temporary larval sites, such as the seasonal backwaters typical of the Malian Sudanese savanna, favors the *Anopheles gambiae* complex, the main vector of *Wuchereria bancrofti* lymphatic filariasis [16,17] In Siradjouba, the increased prevalence of permanent larval habitats, particularly polluted stagnant water in urban and peri-urban areas, supports populations of *Culex spp*., confirmed vectors of arboviruses such as West Nile fever and chikungunya [18,19] (Table 1).

The drastic decline in *Anopheles* in 2023 (1.3% of total captures, exclusively in Konfra and completely absent in Siradjouba) is likely due to environmental disturbances affecting breeding sites, such as habitat drying or chemical pollution. *Anopheles mosquitoes*, which are highly sensitive to hydrological fluctuations and water salinity, have limited larval plasticity compared to *Culex* mosquitoes [16,20,21] (table 1).

The genus *Culex spp*. retained its absolute predominance, with 3,512 individuals in 2022 compared to 231 in 2023, and a stable distribution favoring Siradjouba (approximately 63-64%). This resilience, despite interannual variations, reflects the high adaptability of *Culex* to diverse ecological niches, including organically enriched and pollutant-tolerant waters, which is a major challenge for arbovirus surveillance and control [12,22,23] (Table 1).

The overall reduction in the density of both genera in 2023 suggests interannual climate variability, potentially linked to rainfall patterns, or local public health interventions, such as mass drug administrations (MDAs) of antiparasitic drugs against filariasis in the Sikasso region [24] (Table 1).

### Effectiveness of *Anopheles gambiae* complex capture methods by location and year

The clear predominance of captures in Konfra compared to Siradjouba suggests a marked difference in ecological or anthropogenic conditions between these two localities [25,26]. This disparity can be explained by the availability and quality of breeding sites, human density, or local practices influencing the presence of mosquitoes [27]. The predominance of the *Anopheles gambiae* complex in Konfra is consistent with its known ecology, which favors clear, temporary waters for egg laying, often associated with inhabited or agricultural areas [16,27] (Table 2).

Pyrethrum Spray Catch (PSC) significantly outperforms other methods for the *Anopheles gambiae* complex (91% of 989 specimens, especially in Konfra), while the Ifakara tent captures only 9% and the Gravid Trap (GT) captures none in 2022. In 2023, PSC remains exclusive for the rare captures. This confirms the literature: PSC effectively targets endophagous females resting inside rooms, which is typical behavior for the *Anopheles gambiae* complex in rural Mali [1,3,28]; the Ifakara tent, which attracts females seeking blood, is less sensitive here [29], and the absence of captures with the GT trap indicates that this method is not well suited to this species complex in these contexts, as has also been documented in other studies. GT specifically targets pregnant females attracted to polluted water and is ineffective without nearby larval habitats [2] (Table 2).

These results demonstrate the superior effectiveness of Pyrethrum Spray Catch (PSC) for capturing the *Anopheles gambiae* complex. It remains the optimal method for monitoring the *Anopheles gambiae* complex in this context, particularly in Konfra, while Ifakara underperforms as a backup and GT is useless here, highlighting the importance of active indoor captures.

### Effectiveness of Culex *spp*. capture methods by location and year

For the 3,743 *Culex spp*., GT dominates in 2022 (54%, especially Siradjouba at 79%), followed by PSC (37%, better in Konfra) and Ifakara (10%). In 2023, Ifakara takes the lead (43%, dominant in Siradjouba), with GT and PSC following.

These results show that the GT trap was overall the most productive in 2022, particularly in Siradjouba, while the PSC method yielded better results in Konfra. The Ifakara trap also proved to be more effective in Siradjouba. In 2023, the Ifakara trap became dominant in Siradjouba, while the GT trap yielded more consistent results between the two locations. This variability reflects the ecological and behavioral differences of *Culex spp*. between sites [30]. The success of the GT in Siradjouba suggests high gravid activity, i.e., females searching for oviposition sites, which is consistent with the biology of *Culex spp*., which often lay eggs in stagnant water rich in organic matter [5,30]. The PSC, which is effective in Konfra, targets endophilic mosquitoes that rest in dwellings, indicating a more endophilic behavior of local populations [5,28]. Studies show that gravid female traps are also known to specifically target females ready to lay eggs, which explains their high productivity in certain contexts [5,31]. The Ifakara tent, designed to capture blood-seeking mosquitoes, complements these methods by targeting different behaviors [15] (Table 3).

These results confirm that the effectiveness of different methods is closely linked to local ecology and mosquito behavior. GT traps are a priority in areas where permanent breeding sites are significant (e.g., in Siradjouba in 2022) [32]. PSC remains essential for sampling endophilic populations, particularly if human habitats are favorable [33]. The Ifakara tent is an essential complementary tool in exophilic environments [29]. These observations highlight the importance of a multi-methodological approach to monitoring *Culex spp*., combining gravid traps, PSC, and Ifakara tents to cover different behaviors (biting, endophilic resting, nocturnal activity) [2] (Table 3).

### Variations in the total number of mosquitoes captured in the two villages in 2022 and 2023

The graph shows a clear predominance of *Culex spp*. in Siradjouba, while members of the *Anopheles gambiae* complex are very poorly represented. This suggests that the ecological conditions in Siradjouba (presence of stagnant water rich in organic matter, pollution, latrines, etc.) are particularly favorable to the proliferation of mosquitoes of the genus *Culex spp.*, known for their ecological plasticity and their ability to colonize anthropized or polluted environments [34]. In Konfra, the distribution between the two genera is more balanced. This indicates the coexistence of larval sites suitable for both groups: clear, temporary waters for *Anopheles* [35,36] and stagnant or polluted waters for *Culex spp.* [34]. However, the population of the *Anopheles gambiae* complex remains larger in Konfra than in Siradjouba (Figure 2). This graph clearly illustrates the ecological variability of mosquito vectors depending on the local context, with major implications for entomological surveillance and vector control. Control strategies must therefore be adapted to take into account the specific composition of mosquito populations and the epidemiological risks associated with each locality (Figure 2).

### Conclusion

The results highlight varying trap performance depending on mosquito species and ecological contexts. For the *Anopheles gambiae* complex, the PSC trap proved to be significantly superior (in Konfra and Siradjouba), while the Ifakara trap showed moderate effectiveness and the gravid trap proved ineffective (at both sites). In contrast, for *Culex spp*., the gravid trap dominates, especially in Siradjouba, and remains competitive in Konfra. The PSC is therefore the preferred method for targeting members of the *Anopheles gambiae* complex in temporary or permanent environments, while the gravid trap excels for *Culex spp*. in permanent sites. The Ifakara trap is a moderate alternative, better suited to *Anopheles gambiae*.

For future xenomonitoring studies, the PSC and the trap for pregnant females should be used in combination to capture mosquitoes, followed by qPCR on pooled mosquito samples to detect *Wuchereria bancrofti* DNA, even at low densities.

## Acknowledgments

I would like to express my sincere thanks to:

- The Coalition for Operational Research on Neglected Tropical Diseases, funded by the Task Force for Global Health, primarily through the Bill & Melinda Gates Foundation, the U.S. Agency for International Development via its Neglected Tropical Diseases program, and the British public for their financial support. The grant was administered by the African Network for Research on Neglected Tropical Diseases (ARNTD). The content of this work is the sole responsibility of [Lamine Soumaoro] and does not necessarily reflect the views of COR-NTD, USAID, UK aid, or ARNTD.
- The U.S. National Institutes of Health (NIH), through Dr. Thomas Nutman of the Section on Helminth Immunology and Parasitic Diseases, National Institute of Allergy and Infectious Diseases.

## Declarations

### Ethical Approval and Consent to Participate

The Ethics Committee (EC) of the University of Science, Engineering, and Technology of Bamako approved the protocol under number 2022/294/EC/USTTB.

As part of the mosquito captures using the three collection methods, the team obtained verbal consent from villagers as well as from individuals working as vector collectors with the research team, and from the owners of the rooms visited for mosquito collection.

All participants were free to withdraw from the study at any time. In accordance with the Declaration of Helsinki, our protocol was submitted to and approved by an ethics committee.

### Consent for publication

During the initial study visits, details of the study’s objectives and implementation phases were clearly explained to the various village chiefs in a series of meetings, so that they would be informed and engaged as beneficiaries of the results, which would help the national program to improve the quality of lymphatic filariasis surveillance to rapidly detect any re-emergence of the disease in their areas bordering the country. Their verbal consent was obtained for:

- The sharing of reports with the head of the national lymphatic filariasis elimination program and the chief medical officer of the two health districts, as well as with the health managers of the health centers in the study area.
- Publication of the final report and articles with all stakeholders (from national to very peripheral levels, using appropriate media).
- Presentations of results at national and international conferences and relevant meetings to share results widely for better use and impact.

### Competing interests

The authors declare no competing interests.

### Clinical trial

Not applicable.

### Funding

This publication was supported by the Coalition for Operational Research on Neglected Tropical Diseases, which is funded by the Task Force for Global Health, principally by the Bill & Melinda Gates Foundation, the US Agency for International Development through its Neglected Tropical Diseases Program, and with assistance from the people of the UK. The grant was administered by the African Research Network on Neglected Tropical Diseases (ARNTD). The contents are the responsibility of [Lamine Soumaoro] and do not necessarily reflect the views of COR-NTD, USAID, British Aid or ARNTD. Sponsored Research Agreement (Ref: SGPV/0310.358). The study was funded in part (for TBN) by the Division of Intramural Research (DIR) of the National Institute of Allergy and Infectious Diseases, NIAD, providing us with the primers for the PCR reactions.

### Data availability

The data used in this manuscript can be found through the International Center of Excellence in Research of the University of Science, Techniques and Technologies of Bamako by sending an email to the principal investigator and first author of the manuscript.

### Author contributions

LS wrote this article and created the illustrations, and MEC, AAD, MS, AFD, SYC, SSD, MMK, AKK, KD, HD, ASY, and YIC have all read and corrected the article.

## Notes

### Competing Interest Statement

The authors have declared no competing interest.

